# Brequinar and Dipyridamole in Combination Exhibits Synergistic Antiviral Activity Against SARS-CoV-2 *in vitro*: Rationale for a host-acting antiviral treatment strategy for COVID-19

**DOI:** 10.1101/2022.03.30.486499

**Authors:** James F. Demarest, Maryline Kienle, RuthMabel Boytz, Mary Ayres, Eun Jung Kim, Donghoon Chung, Varsha Gandhi, Robert Davey, David B. Sykes, Nadim Shohdy, John C. Pottage, Vikram S. Kumar

## Abstract

The continued evolution of severe acute respiratory syndrome coronavirus 2 (SARS-CoV-2) has compromised the efficacy of currently available vaccines and monoclonal antibody (mAb)-based treatment options for COVID-19. The limited number of authorized small-molecule direct-acting antivirals present challenges with pill burden, the necessity for intravenous administration or potential drug interactions. There remains an unmet medical need for effective and convenient treatment options for SARS-CoV-2 infection. SARS-CoV-2 is an RNA virus that depends on host intracellular ribonucleotide pools for its replication. Dihydroorotate dehydrogenase (DHODH) is a ubiquitous host enzyme that is required for de novo pyrimidine synthesis. The inhibition of DHODH leads to a depletion of intracellular pyrimidines, thereby impacting viral replication in vitro. Brequinar (BRQ) is an orally available, selective, and potent low nanomolar inhibitor of human DHODH that has been shown to exhibit broad spectrum inhibition of RNA virus replication. However, host cell nucleotide salvage pathways can maintain intracellular pyrimidine levels and compensate for BRQ-mediated DHODH inhibition. In this report, we show that the combination of BRQ and the salvage pathway inhibitor dipyridamole (DPY) exhibits strong synergistic antiviral activity in vitro against SARS-CoV-2 by enhanced depletion of the cellular pyrimidine nucleotide pool. The combination of BRQ and DPY showed antiviral activity against the prototype SARS-CoV-2 as well as the Beta (B.1.351) and Delta (B.1.617.2) variants. These data support the continued evaluation of the combination of BRQ and DPY as a broad-spectrum, host-acting antiviral strategy to treat SARS-CoV-2 and potentially other RNA virus infections.

## INTRODUCTION

As of March 2022, there have been more than 6-million reported deaths and greater than 400-million cases of coronavirus disease 2019 (COVID-19) worldwide as suggested by the World Health Organization (WHO; https://covid19.who.int/). Furthermore, the number of infections with severe acute respiratory syndrome coronavirus 2 (SARS-CoV-2), the causative agent of COVID-19, likely exceeds the number of reported cases with an estimated excess mortality of 18.2 million [1]. Despite the availability of multiple prophylactic vaccines, SARS-CoV-2 continues to evolve, compromising the efficacy of vaccines and monoclonal antibody (mAb)-based treatment options. Furthermore, there are only a handful of small-molecule antivirals including two RNA-dependent RNA polymerase (RdRp) inhibitors, remdesivir and molnupiravir, and one inhibitor of the SARS-CoV-2 main protease (Mpro), nirmatrelvir (nirmatrelvir also requires the pharmacologic booster ritonavir in order to achieve sufficient plasma drug levels). There remains therefore a high unmet medical need for safe, efficacious, and patient-friendly treatments for SARS-CoV-2 infection.

While the therapeutic small molecules described above are direct-acting antivirals (DAAs) that target virus-specific proteins, an alternative treatment strategy is to develop host-acting antivirals (HAAs) that target host pathways essential for the viral lifecycle. A host-based mechanism of action may provide unique advantages over DAAs including a greater likelihood of broad-spectrum antiviral activity against several families of RNA viruses. In addition, HAAs may possess an inherently higher barrier to the development of resistance when compared to DAAs as host-targets typically remain unchanged in contrast to the rapid emergence of viral variants containing mutations that decrease the efficacy of DAAs.

Dihydroorotate dehydrogenase (DHODH) is a host enzyme that is essential for *de novo* pyrimidine synthesis and has emerged as a candidate target of HAAs. Brequinar (BRQ) is an orally available, selective, and potent low nanomolar human DHODH inhibitor (DHODHI) shown to deplete intracellular uridine, cytidine, and thymidine levels *in vitro* and *in vivo* [2, 3, 4]. As an antiviral approach, DHODHIs such as BRQ block the host production of cellular pyrimidine nucleotide triphosphates (NTPs) required by viruses for replication [5, 6, 7, 8]. To date, BRQ has been studied in multiple human clinical studies that have included more than 1000 subjects with viral infection, hematologic malignancies, or autoimmune disorders [Clear Creek Bio data on file; 2, 3, 9, 10].

DHODHIs exhibit potent *in vitro* activity against several RNA viruses such as SARS-CoV-2, influenza, Zika (ZKV), Dengue (DENV), respiratory syncytial virus (RSV), and Ebola (EBOV) [5, 6, 7, 8]. Despite this promising *in vitro* data, the successful translation of DHODHI monotherapy showing clinical benefit is lacking. While the DHODHI NITD-982 exhibited *in vitro* antiviral activity against DENV, the *in vivo* treatment of infected mice had no effect on viremia [11]. This is likely due to other pathways that can compensate for the reduction of pyrimidine NTPs by DHODH inhibition such as the host nucleoside transporters (equilibrative nucleoside transporters, ENT) that facilitate salvage via extracellular uridine and cytidine [12, 13]. The concentration of extracellular uridine in mammalian plasma may range up to 10 μM [12] and may have contributed to limited clinical efficacy of DHODH inhibition in the oncology setting [2].

A combination of DHODH and nucleotide salvage pathway inhibitors may therefore be required for optimal therapeutic efficacy (**Figure 1**). Consistent with this hypothesis, the antiviral activity of a DHODHI against DENV was enhanced by the addition of cyclopentenyl uracil, an inhibitor of uridine salvage [14]. BRQ has also been shown to synergize with the ENT1/2/4 inhibitor dipyridamole (DPY) which blocks the transport of extracellular pyrimidines needed for *in vitro* growth of tumor cell lines [13]. DPY, FDA approved in 1961 [https://www.accessdata.fda.gov/drugsatfda_docs/label/2019/012836s061lbl.pdf], also inhibits platelet aggregation and has widespread clinical use in combination with aspirin for the secondary prevention of stroke [https://www.accessdata.fda.gov/drugsatfda_docs/label/2012/020884s030lbl.pdf]. We chose to evaluate the *in vitro* pharmacologic and antiviral activity of the combination of BRQ and DPY (BRQ+DPY) in uninfected and SARS-CoV-2-infected A549/ACE2 cells.

**Figure 1:**
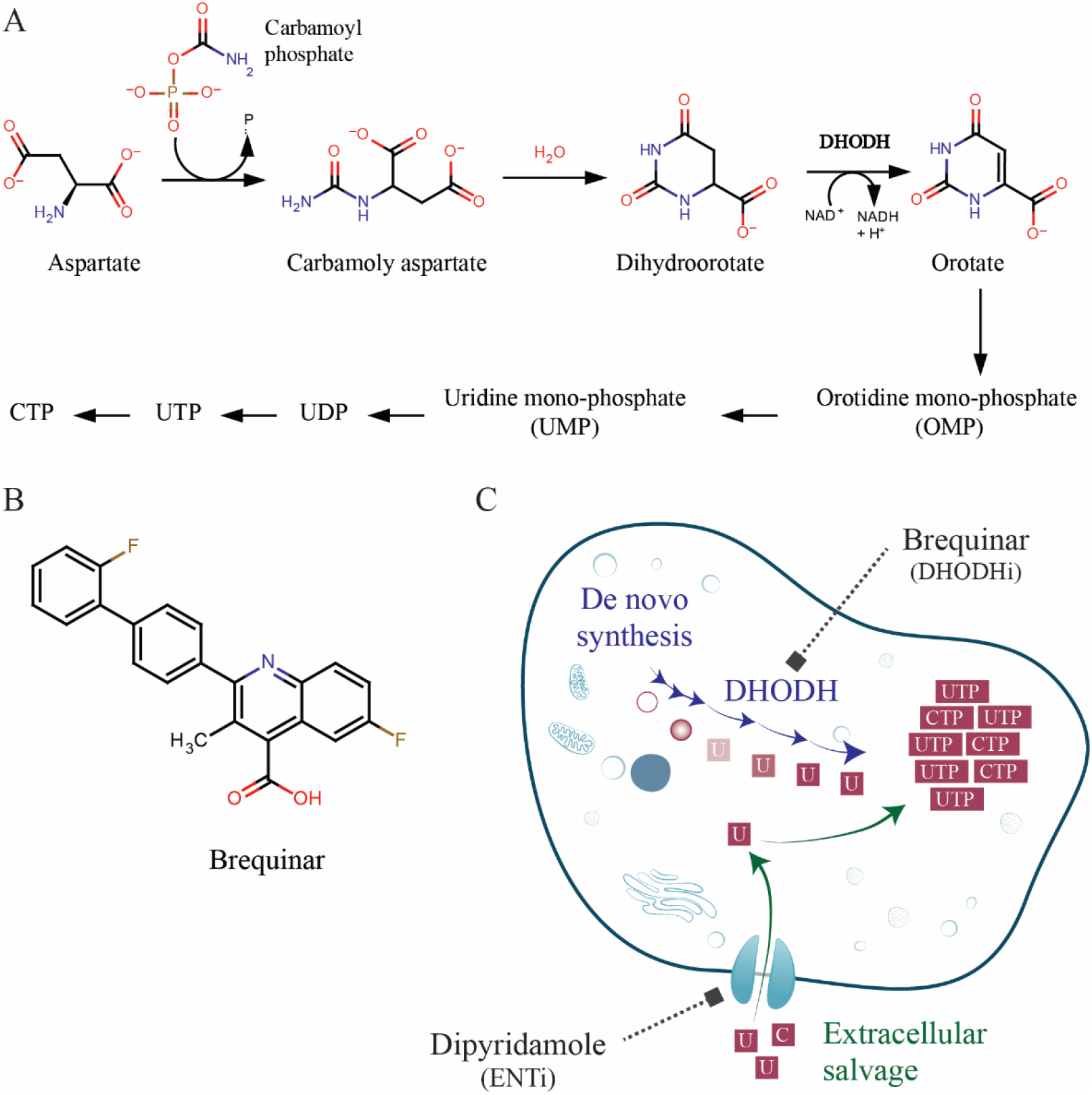
Rationale for combined DHODH and salvage pathway inhibition

## RESULTS

### DPY potentiates the suppression of pyrimidine NTPs by BRQ without apparent cytotoxicity

To study the effect of BRQ+DPY on the concentration of cellular pyrimidine nucleotides (CTP and UTP), uninfected A549/ACE2 cells were treated either with BRQ, DPY, or in combination, over an 8-hour time course. The A549/ACE2 cell line was derived from the parental lung carcinoma A549 and engineered to express human ACE2, the entry receptor for SARS-CoV-2, for use in antiviral assays. Cells were harvested after treatment at each time point (0, 1, 2, 4, and 8 hours) and the NTP concentrations were compared to the control group (T=0, DMSO treated) (**Figure 2**).

**Figure 2.**
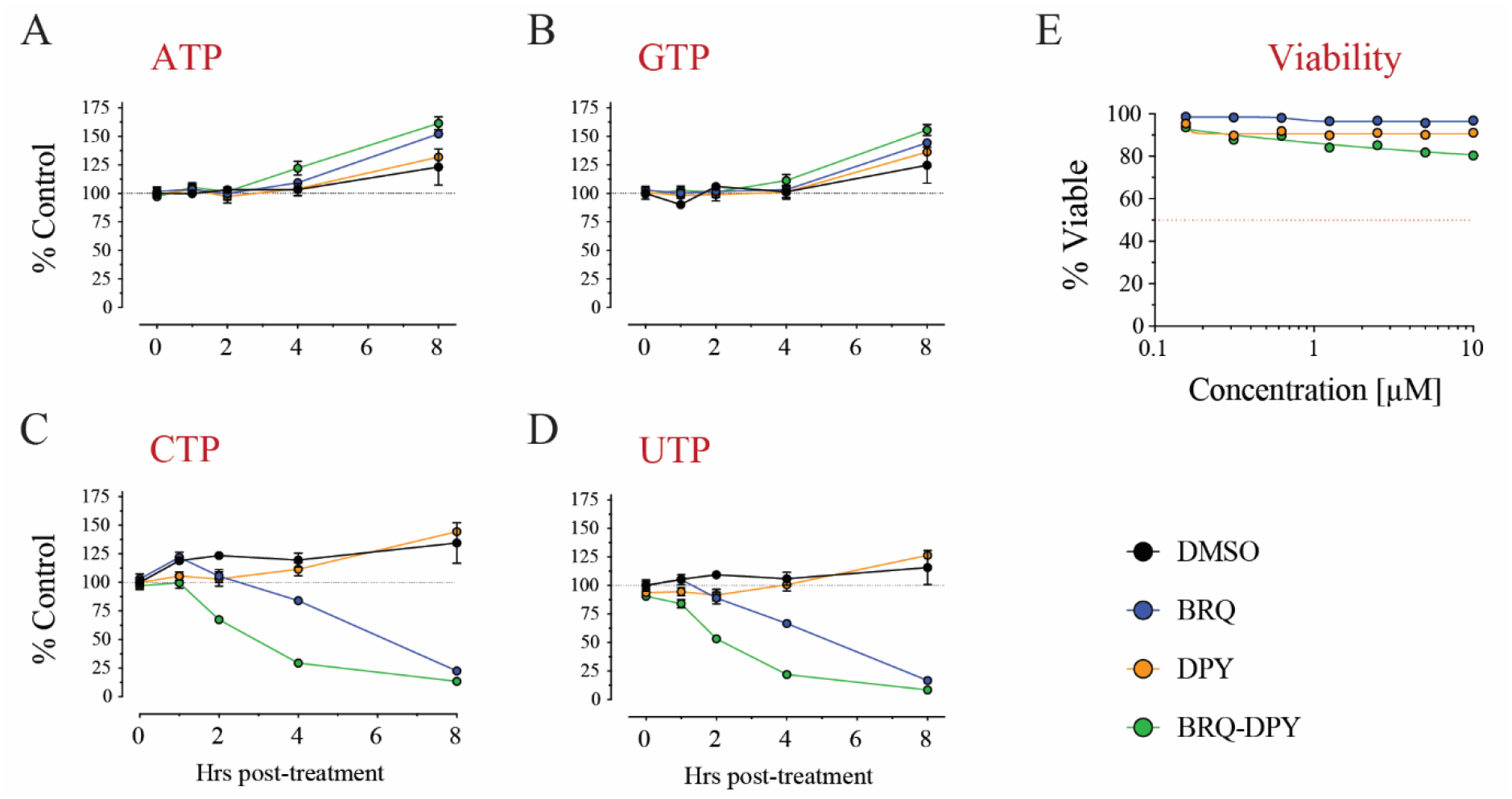
DPY potentiates the suppression of pyrimidine NTPs by BRQ without cytotoxicity. **a-d)** Changes in cellular nucleotide pools over 8 hours in response to the treatment of brequinar (BRQ; 1 μM) and dipyridamole (DPY; 1 μM) for single and combination treatment. Free NTP concentrations were normalized to the vehicle control group at 0 hours post treatment (n=3 per time point, per group). **e)** A549/ACE2 cell viability following treatment of BRQ and DPY for three days. Cell viability was measured with CellTiter-Glo and compared to the vehicle control group (DMSO treated).

The concentrations of purine nucleotides (ATP and GTP) were not decreased by any of the treatments. As a single agent, DPY (1 μM) had no impact on either UTP or CTP levels relative to control at time 0 (baseline). Single agent BRQ (1 μM) exhibited a reduction in pyrimidine NTPs (CTP and UTP) concentrations starting at 4 h (16% and 33% decrease for CTP and UTP, respectively) and the effect was more pronounced at 8 h (77.5% and 83.32% decrease for CTP and UTP, respectively) (**Figure 2).** The combination of BRQ (1 μM) + DPY (1 μM) demonstrated a potentiated effect compared to either agent alone. At 2 h, BRQ+DPY reduced CTP and UTP from baseline by 33.6% and 47.3%, respectively. At 4 h, BRQ+DPY reduced CTP and UTP concentrations from baseline by 71% and 79%, respectively; this was similar to what was seen with BRQ alone at 8 h. At 8h, BRQ+DPY reduced CTP and UTP concentrations from baseline by 86.6% and 91.6%, respectively. The effect of BRQ+DPY on pyrimidine NTP levels was significant relative to DMSO controls and BRQ as a single agent (Supplementary Tables).

We queried whether the effect of BRQ+DPY on pyrimidine NTP concentrations was driven by general cytotoxicity. The CC_50_ values for BRQ, DPY, or BRQ+DPY (1:1 ratio) were >10 μM even after three days of exposure. Notably, these concentrations of BRQ and DPY are 10-fold higher than those used to assay the inhibition of pyrimidine biosynthesis and nucleoside salvage, respectively.

### BRQ+DPY suppresses pyrimidine nucleotides even with high exogenous uridine

Given the mechanism of action as a nucleoside transport inhibitor, we asked whether DPY could potentiate the BRQ-mediated decrease in concentration of pyrimidine NTPs even in the presence of higher concentrations of exogenous uridine. To address this, we repeated the experiment in the presence or absence of uridine (20 μM) at 4 h post compound addition in HEK-293T-hACE2 (**Figure 3A**) and A549/ACE2 cells (**Figure 3B**). Pyrimidine NTP levels were substantially reduced with single agent BRQ (1uM) in both cell lines: HEK-293T-hACE2 (88% and 94% decrease for CTP and UTP, respectively) and A549/ACE2 (56% and 63% decrease for CTP and UTP, respectively). The combination of BRQ (1 μM) and DPY (1 μM) exhibited greater reduction of pyrimidine NTP levels than BRQ alone in both HEK-293T-hACE2 (87% and 96% decrease for CTP and UTP, respectively) and A549/ACE2 (91% and 94 % for CTP and UTP, respectively) cells; the latter being consistent with our previous experiment (**Figure 2**).

**Figure 3:**
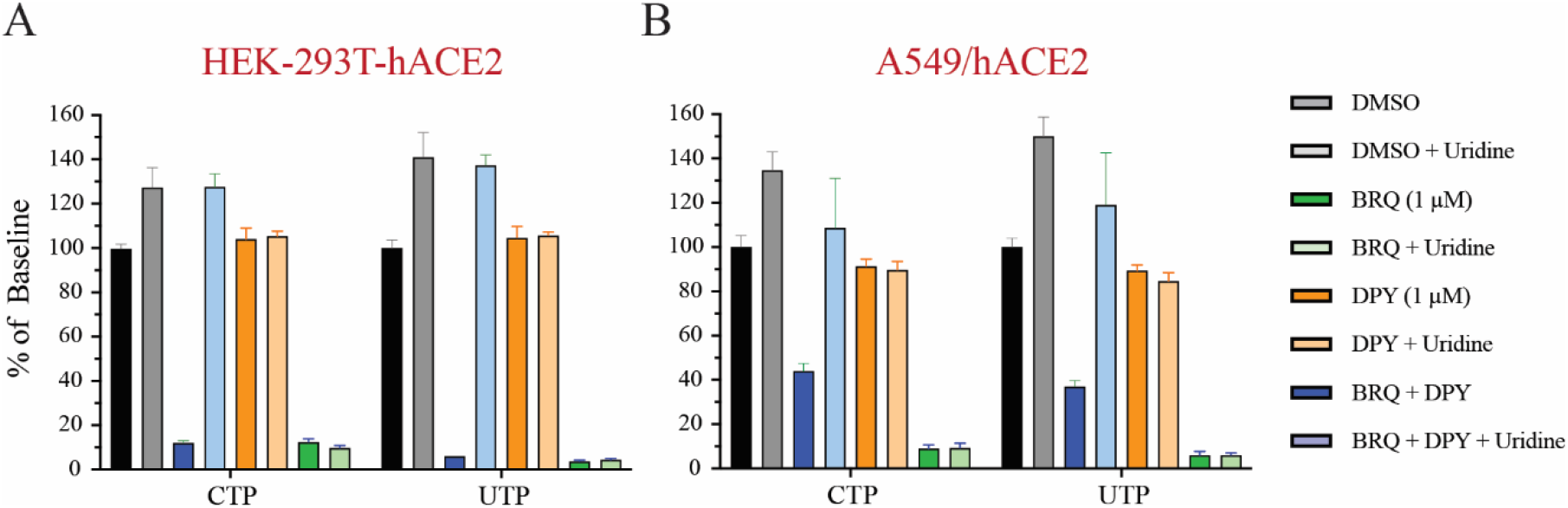
BRQ+DPY suppresses levels of pyrimidine nucleotides even in presence of high concentrations of exogenous uridine. The ability of 1 μM BRQ or 1 μM DPY alone or in combination to decrease intracellular pyrimidine nucleotides concentrations in HEK-293T-hACE2 (Panel A) and A549/ACE2 (Panel B) at 4 h post compound addition; the effect of BRQ alone was more pronounced in HEK-293T-hACE2 cells. The addition of excess exogenous uridine (20 M) was able to rescue pyrimidine nucleotides inhibition by BRQ alone but not by the combination BRQ+DPY in both HEK-293T-hACE2 and A549-ACE2 cells. Purine nucleotides levels were unaffected in either cell type (data not shown).

The addition of excess uridine rescued cellular CTP and UTP concentrations in cells treated only with BRQ. In contrast, and consistent with the pharmacological action of DPY, rescue of CTP and UTP levels with excess uridine was not observed in cells treated with BRQ+DPY. Purine NTP levels were not impacted by BRQ or DPY or the combination (data not shown). This result confirms that blocking pyrimidine salvage potentiates the BRQ effect on intracellular pyrimidine NTP concentrations in cells even in the presence of excess concentrations of extracellular uridine.

### BRQ+DPY exhibits synergistic antiviral activity *in vitro* against SARS-CoV-2

Given their mechanisms of action and our *in vitro* data showing that the combination of BRQ+DPY rapidly lowers pyrimidine NTP pools which are required for viral RNA synthesis/replication (**Figure 2**), we explored whether BRQ+DPY may also exhibit greater antiviral activity than either agent alone.

#### BRQ+DPY demonstrates synergistic antiviral activity

The antiviral activity of BRQ+DPY inhibition was assessed in A549/ACE2 cells infected with SARS-CoV-2 Beta variant (B.1.351) (**Figure 4**). DPY treatment alone had no impact on SARS-CoV-2 infection (data not shown). The antiviral activity of BRQ was enhanced in a dose-dependent manner by the addition of DPY (**Figure 4A** and **Table 1**). The decrease in EC_50_ values was observed across pharmacologically relevant concentrations of DPY (0.78 μM or 1.563 μM; [15]); decreases from 2.67 μM for BRQ alone to 0.80 μM (3.3-fold lower) and to 0.59 μM (4.5-fold lower), respectively. Similar enhancement of BRQ antiviral activity with the addition of DPY was also observed in separate experiments using Vero cells infected with geographically distinct Wuhan-related SARS-CoV-2 strains (data not shown). The antiviral activity of BRQ+DPY was not attributable to cellular cytotoxicity as cytotoxicity of BRQ and DPY alone or as a combination of BRQ (5.0 μM max) and either DPY at 0.78 μM or 12.5 μM was <50% in the MTT assay (Supplemental Figure S1).

**Figure 4:**
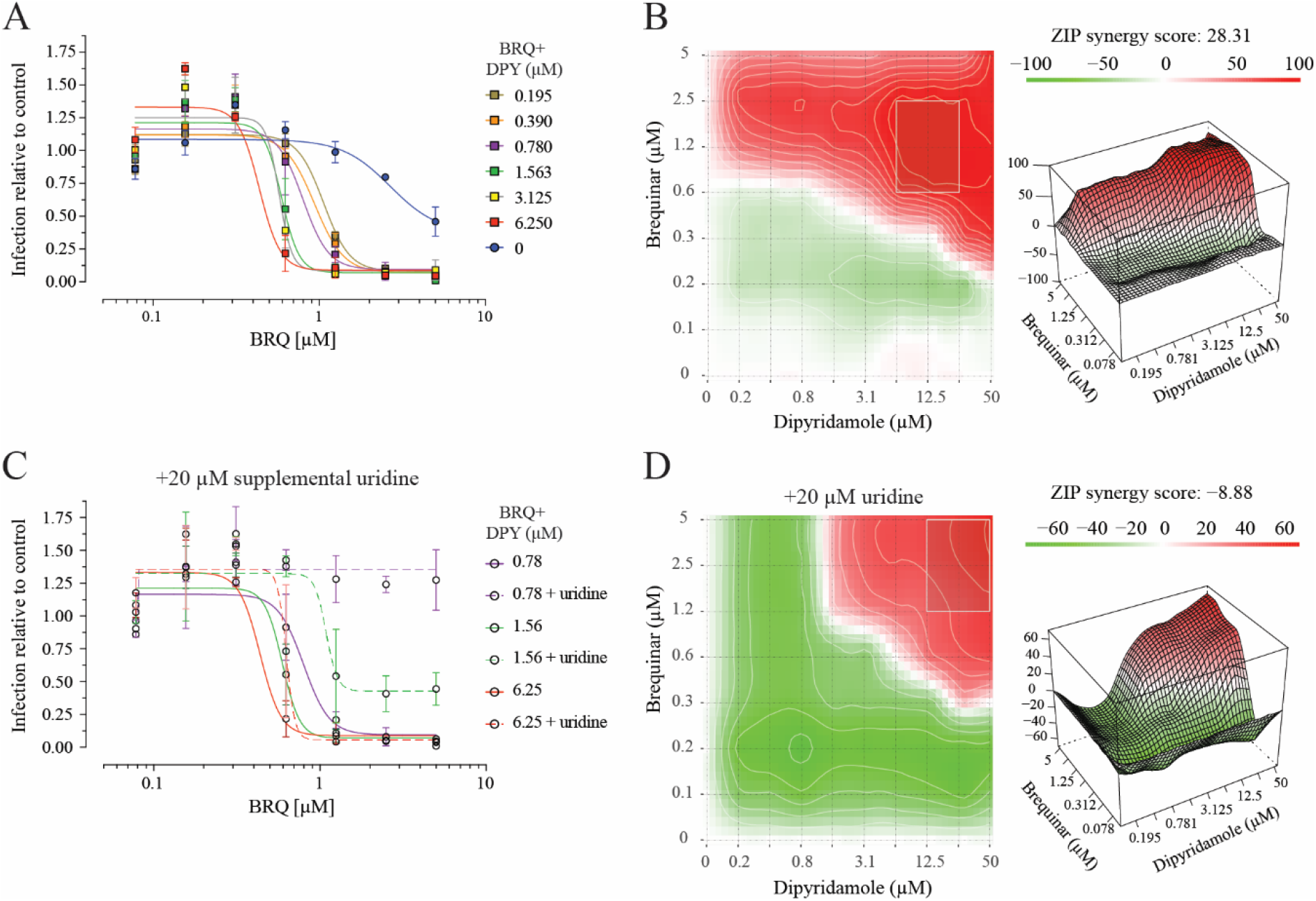
BRQ+DPY exhibits strong synergistic antiviral activity that is partially abrogated by supplementation of extracellular uridine. A) The antiviral activity of BRQ alone was enhanced in a dose-dependent manner by the addition of DPY. BRQ at concentrations of 0.078 – 5.0 μM alone or in combination with DPY at concentrations of 0.195 – 50.0 μM were evaluated. B) The 2-drug combination of BRQ+DPY exhibited strong synergy with highest effect observed at BRQ 0.6 – 2.5 μM and DPY 6.2 – 25 μM. C) The addition of excess exogenous uridine reduces the dose-dependent effect of BRQ+DPY. D) Addition of 20 μM uridine shifts the area of greatest synergy towards higher concentrations of both BRQ and DPY, and synergy scores suggest that uridine addition abrogates synergy still observed at higher DPY concentrations, rendering the effect of the drug combination overall as additive. A549-ACE2 cells were dosed with combinations of drugs and challenged with SARS-CoV-2 infection for 48 h. Infection was quantified from IFA images stained for SARS-CoV-2 N protein, and synergy was calculated using SynergyFinder2.0 [16] under the LL4 curve and ZIP models. Dark red areas and peaks in the contour plot indicate strong synergistic interaction, with the white box representing the area of strongest synergy. Synergy scores greater than 10 indicate a synergistic interaction, −10 to 10 suggests additivity, and less than −10 supports antagonism.

**Table 1.** BRQ+DPY Synergy vs SARS-CoV-2 Beta (B.1.351)

| Treatment | EC<sub>50</sub> [uM]<br>No Added Uridine | Fold-Change<br>vs<br>BRQ Alone | EC<sub>50</sub> [uM]<br>With 20uM Uridine | Fold-Change<br>vs<br>BRQ Without<br>Added Uridine |
| --- | --- | --- | --- | --- |
| DPY Alone | >50 | N/A | >50 | N/A |
| BRQ Alone | 2.67 | 1.00 | >5 | >1.87 |
| BRQ + 0.195uM DPY | 1.06 | -2.51 | >5 | >4.70 |
| BRQ + 0.390uM DPY | 0.94 | -2.85 | >5 | >5.34 |
| BRQ + 0.789uM DPY | 0.80 | -3.35 | >5 | >6.27 |
| BRQ + 1.563uM DPY | 0.59 | -4.50 | 1.09 | 1.84 |
| BRQ + 3.125uM DPY | 0.57 | -4.66 | 0.90 | 1.57 |
| BRQ + 6.250uM DPY | 0.44 | -6.12 | 0.63 | 1.44 |
| BRQ + 12.5uM DPY | 0.32 | -8.24 | 0.40 | 1.23 |
| BRQ + 25uM DPY | 0.28 | -9.48 | 0.28 | 1.00 |
| BRQ + 50uM DPY | 0.28 | -9.53 | 0.27 | 1.00 |
| **Synergy** |  |  |  |  |
| Synergy Score (Ave)* | 22.57 | Synergy | -4.66 | Additivity |
| Most Synergistic Area | 75.15 |  | 58.36 |  |

\*Determined with ZIP (Bliss+Loewe); Synergy scores: >10.0 indicates synergy; -10 to 10 indicates additivity; < -10 indicates antagonism; N/A: Not Applicable.

To determine whether BRQ+DPY had additive, synergistic, or antagonistic interactions, the antiviral data were analyzed using a combination of Loewe and Bliss models [16]. The 2-drug combination of BRQ+DPY exhibited a strong synergistic interaction (**Figure 4B** and **Table 1**). The average synergy score from three replicates was 22.6, with the most synergistic area covering 75.2 arbitrary square units. These data confirm synergistic antiviral activity of BRQ+DPY in the treatment of SARS-CoV-2 at pharmacologically relevant drug concentrations.

#### Exogenous uridine supplementation partially abrogates the synergistic effect

The effect of excess exogenous uridine on the antiviral activity of BRQ+DPY was more pronounced at DPY concentrations <3 μM where substantial or complete reversal of antiviral activity (e.g., EC_50_ > maximum concentration tested) was observed. DPY concentrations at or above 6.25 μM decreased the impact of excess uridine (**Table 1, examples in Figure 4C**). Consistent with the increase in EC_50_ values for BRQ+DPY in the presence of 20 μM uridine, synergy analysis of infection data revealed a shift in synergistic dose combinations to higher concentrations of both drugs (**Figure 4D**). Furthermore, the synergy score in the presence of excess exogenous uridine was −4.7, which falls in the range of additivity. The additive score is in part confounded by the apparent antagonism of excess uridine at low concentrations of DPY and by the lesser effect at higher DPY concentrations. The physiological relevance of 20 μM uridine, some 2- to 9-fold higher than physiologic levels [12], is not clear.

#### BRQ antiviral activity against SARS-CoV-2 Delta Variant of Concern (B.1.617.2) is enhanced with low concentrations of DPY

Based on the findings with SARS-CoV-2 Beta (B.1.351), the antiviral activity of BRQ+DPY was evaluated in the same assay system using the SARS-CoV-2 Delta variant of concern (VOC) (B.1.617.2) (**Figure 5**). BRQ+DPY also exhibited enhanced antiviral activity against SARS-CoV-2 Delta (B.1.617.2), relative to DMSO control as well as single agent BRQ.

**Figure 5:**
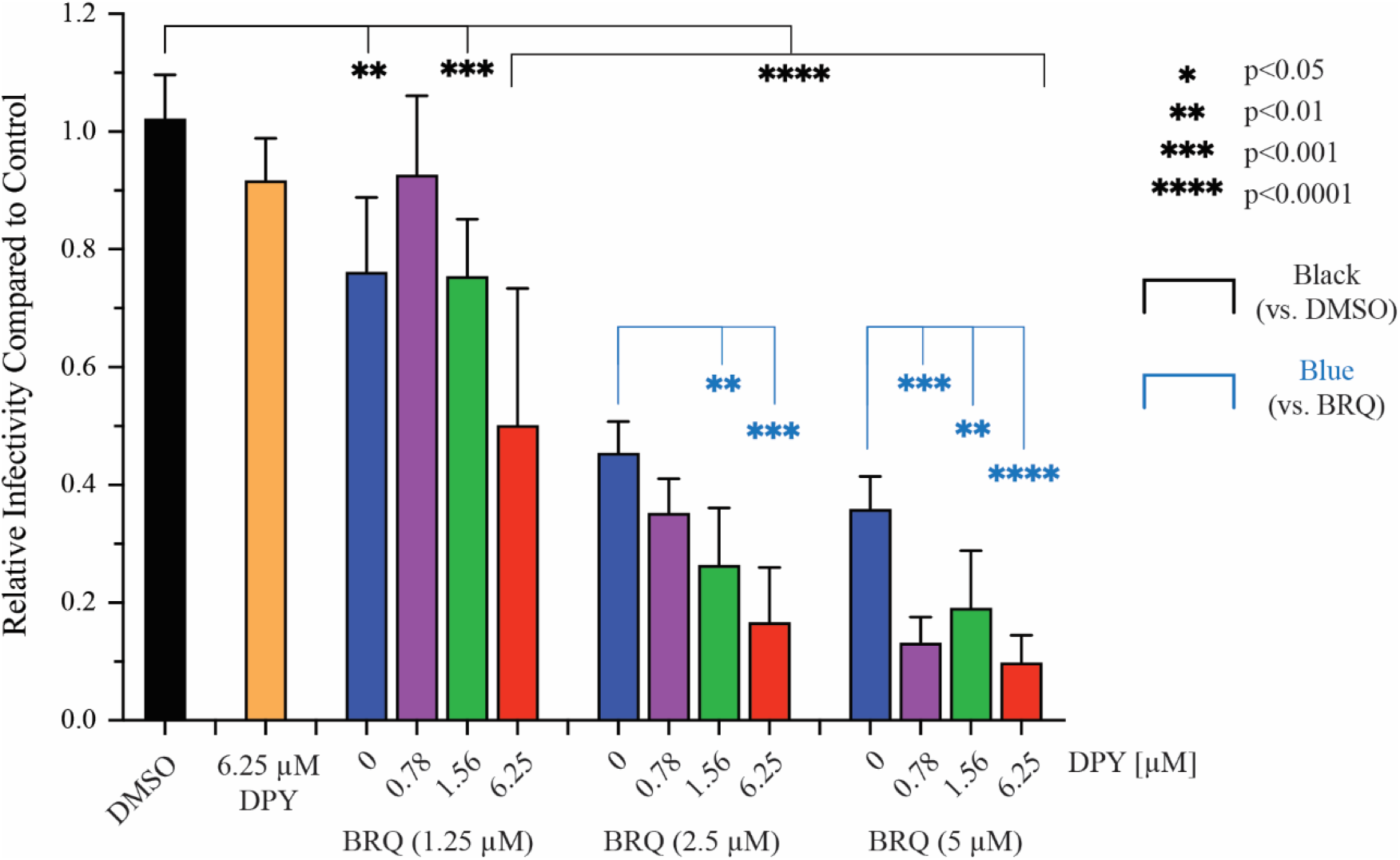
BRQ antiviral activity against SARS-CoV-2 Delta VOC (B.1.617.2) is enhanced with low concentrations of DPY. The antiviral activity of BRQ alone was enhanced in a dose-dependent manner by the addition of low concentrations of DPY. BRQ at concentrations of 1.25, 2.5 and 5.0 μM alone or in combination with DPY at concentrations of 0.78, 1.56 and 6.25 μM were evaluated. A549-ACE2 cells were dosed with combinations of drugs and challenged with SARS-CoV-2 infection for 48 h. Infection was quantified from IFA images stained for SARS-CoV-2 N protein.

## DISCUSSION

There remains a significant unmet medical need for safe and convenient treatments for people infected with SARS-CoV-2. The most recently authorized antivirals have challenges with pill burden, inconvenient routes of administration, or potential drug interactions. The potential efficacy of DHODHI monotherapy for treatment of viral infection in humans remains to be proven. Clinical experience in the oncology space suggests that the impact of available extracellular uridine and the activity of nucleoside transporters may limit the efficacy of DHODHI monotherapy. The *in vitro* antiviral activity of BRQ alone against SARS-CoV-2 has been reported by others [17, 18, 19]. Given the growing evidence suggesting that DHODH inhibition alone *in vitro* may not translate to *in vivo* activity, we explored a host-target antiviral strategy with the combination BRQ and DPY.

The addition of DPY enhanced the inhibitory effect of BRQ on the concentration of intracellular pyrimidine NTPs *in vitro*. This effect was consistent in the HEK-293T-hACE2 and A549/ACE2 cell lines (**Figs 2-3**) as well as multiple other cell lines and primary cells (data not shown). The depletion of pyrimidine NTP pools by BRQ could be overcome with the addition of excess exogenous uridine (20 μM) whereas the inhibitory activity of BRQ+DPY was preserved in the setting of high concentrations of extracellular uridine. Furthermore, the inhibitory activity associated with low concentrations of BRQ and DPY was not driven by apparent cytotoxicity.

Within a cell infected with SARS-CoV-2, more than 50% of RNA transcripts are viral RNA transcripts, and all nucleotides are host-derived [20]. Thus, potent reduction of pyrimidine NTP pools by the combination of BRQ+DPY should effectively limit viral replication. In A549/ACE2 cells (**Figure 4**), DPY alone had no evidence of antiviral effect against SARS-CoV-2 Beta (B.1.351) and BRQ had marginal single agent activity consistent with published reports [17, 18, 19] and Clear Creek Bio data (not shown). Consistent with the enhanced *in vitro* reduction of pyrimidine NTP levels in uninfected cells with 1 μM BRQ (**Figure 2**), the addition of DPY increased the antiviral activity of BRQ alone in a dose-dependent manner (**Figure 4A** and **Table 1**). It is worth noting that as with the *in vitro* analyses of pyrimidine NTP levels, the antiviral activity of BRQ+DPY was evident at concentrations with potential pharmacologic relevance, <2 μM each [Clear Creek Bio data on file; 9, 15] and was not driven by apparent cytotoxicity even at high concentrations. The analysis of 2-drug antagonism or additivity/synergy using a combination of Loewe and Bliss models demonstrated strong synergy for the combination of BRQ+DPY.

The addition of excess exogenous uridine abrogated the antiviral activity of BRQ in combination with lower concentrations of DPY in SARS-CoV-2 infected A549/ACE2 cells though this abrogation was less pronounced at the higher DPY concentrations (**Figure 4C and Table 1**). The synergy analysis in the presence of 20 μM uridine shifted from synergy to additivity. These data highlight the importance of extracellular uridine in maintaining cellular pyrimidine NTP levels and supports the concept that inhibiting both *de novo* and salvage pyrimidine pathways with BRQ+DPY merits further exploration.

A major area of concern is the development of viral resistance, as seen with selective pressures by DAAs, or viral escape from immune pressures that lead to the emergence of novel SARS-CoV-2 VOCs, as has been observed after vaccination and the development of natural immunity. As BRQ and DPY target host rather than viral proteins, the antiviral activity of BRQ+DPY should have limited liability with respect to viral escape or development of resistance. Our data demonstrated comparable antiviral activity of BRQ+DPY against SARS-CoV-2 strains including those like the original Wuhan-1 as well as Beta and Delta variants of concern.

Given the *in vitro* synergy against multiple SARS-CoV-2 strains in different cell types, a similar enhancement of BRQ antiviral activity may be observed *in vivo* with the host-based combination treatment of BRQ+DPY. Furthermore, given this synergy and the inability of excess exogenous uridine to reverse this, the HAA combination of BRQ + DPY may be expected to present a high barrier to the development of clinically relevant resistance relative to DAAs. This needs to be formally assessed in clinical trials.

In this report, we demonstrated that the combination of BRQ+DPY significantly reduces pyrimidine NTP levels which translates to synergistic antiviral activity against SARS-CoV-2 variants *in vitro*. The antiviral activity observed with BRQ+DPY at pharmacologically relevant concentrations supports continued investigation of this combination as an oral treatment approach for COVID-19. A small, outpatient Phase 2 clinical trial is currently underway to evaluate this concept. [https://clinicaltrials.gov/ct2/show/NCT05166876]. Finally, if this approach is successful, investigation of the use of BRQ+DPY in other RNA viral infections may be warranted.

## MATERIALS AND METHODS

### Test Articles

Brequinar (Selleck Chemicals, Cat# S6626) and dipyridamole (Sigma Aldrich Prod# D9766) were provided by Clear Creek Bio. Uridine was obtained from Sigma (Cat#: U3003-5G) and Alfa Aesar/Fisher (AAA1522706) for the NTP and antiviral assays, respectively.

### Cell lines and virus

A549/ACE2 and HEK-293T-hACE2 (BEI resources, NR-53821 and NR-52511) cells were cultured in DMEM media supplemented with 10% FBS and 2 mM L-glutamine. Cells were passed twice in a week and maintained at 37° C with 5 % CO_2_. The absence of mycoplasma contamination was validated regularly with a PCR-based method (ATCC, Universal Mycoplasma Detection Kit, 30-1012K).

### *In vitro* determination of BRQ+DPY effect on pyrimidine NTPs

Cells were plated in 6-well plates (200,000 cells/well) one day prior to experiment. Test compounds were dissolved in DMSO as a stock and then diluted in culture media before testing. The final concentration of DMSO was kept at 0.25%. For drug treatment, cell supernatant was removed and replaced with media containing test compounds (n=3 per group). At timepoints denoted, cell lysates were prepared according to a protocol published [21] with minor modifications. In brief, cells were washed with PBS and treated with 0.75 mL of with 0.4 N perchloric acid (PCA) on ice and harvested. Extracts were centrifuged (1500 rpm) and supernatants were combined with a second 0.25 ml extraction. Extracts were combined and neutralized with 10 N and 1 N KOH. Neutralization was determined using pH paper. Samples were stored at −20°C until. The NTP concentrations of samples were analyzed with HPLC.

PCA extracts were analyzed using either a Waters 2695e HPLC with a Waters 2489 UV/Visible detector or a Waters 2695 HPLC with a Waters 2487 Dual λ Absorbance detector. A Partisil 10 SAX column separated nucleoside triphosphates at a flow rate of 1.5 ml/min with a 50-minute concave gradient curve (curve 8) from 60% 0.005 M NH4H2PO4 (pH 2.8) and 40% 0.75 M NH4H2PO4 (pH 3.8) to 100% 0.75 M NH4H2PO4 (pH 3.8). Standard ribonucleotides were used to create a standard curve, which was used to quantitate nucleotide pools [21].

Analysis of Variance with Dunnett’s Multiple Comparisons Test was used to test for significant differences from the DMSO Control at Time 0 vs all other conditions or DMSO vs BRQ, DPY, or BRQ+DPY treatment at each timepoint (Supplemental Tables S1 and S2).

For assessment of potential cytotoxicity, cells were seeded in 96-well white plates at a density of 12,000 cells per well and cultured overnight. The next day, test compound diluted in cell culture media were added to the cells at the final concentration from 50 μM to 0.4 μM, 8 points by 2-fold serial dilution. After three days of incubation, cell viability was evaluated with CellTiter-Glo, measuring luminescence with Synergy 4 (Biotek). 1% of Triton-X100 and 0.25% DMSO were used as the positive and negative control, 0 and 100% cell viability, respectively.

### BRQ+DPY antiviral experiments

A549/ACE2 cells were plated in 96-well plates at a density of 10,000 cells per well in RPMI supplemented with 10% FBS and allowed to adhere overnight. The following day cells were treated with 2-fold dilution series in triplicate of brequinar (ranging 5 μM – 0.0781 μM) and dipyridamole (ranging 50 μM – 0.195 μM) in a matrix format allowing each concentration pair to be tested. Dilution series of each compound alone were included on each plate, as well as 10 μM remdesivir positive control and DMSO negative control wells. Exogenous uridine (20 μM) was added to an additional three plates to test whether brequinar inhibition of DHODH-catalyzed uridine synthesis can be overcome by addition of excess uridine. Following one hour of incubation with compounds, cells were challenged with approximately 400 FFU of SARS-CoV-2 Beta (B.1.351; hCoV-19/USA/MD-HP01542/2021) and incubated at 37 ⁰C for 48 h. In a separate experiment using A549/ACE2 cells, dilution series of brequinar and dipyridamole alone or brequinar + 0.78 μM or 12.5 μM dipyridamole were tested with the SARS-CoV-2 Delta VOC (B.1.617.2). Pre-treatment time, infection dose, and length of infection were the same as with Beta variant. Analysis of Variance with Dunnett’s Multiple Comparisons Test was used to test for significant differences from the DMSO Control vs BRQ or BRQ+DPY treatment or between BRQ alone vs BRQ+DPY.

### Immunofluorescence detection of SARS-CoV-2 infection efficiency

Following 48 h of infection plates were fixed in 10% formalin for at least 6 h before removal from the high containment laboratory. Plates were washed in PBS, permeabilized in 0.1% Triton X-100 for 15 min at room temperature and blocked in 3.5% BSA for at least 1 h at room temperature. To detect infection, plates were incubated overnight at 4 ⁰C with a rabbit anti-N protein monoclonal antibody (SioBiological 40143-R004) diluted 1:20,000. Plates were washed and treated with an AF488-conjugated goat anti-rabbit secondary antibody for 2 h at room temperature. Finally, Hoechst33342 was added to visualize cell nuclei. Plates were imaged on a Cytation 1 Multimode Plate Reader (BioTek) using a 4X objective lens. Infection efficiency, defined as GFP-positive cells divided by total nuclei, was calculated for each image using a CellProfiler pipeline.

### Synergy calculations

Drug synergy was calculated with SynergyFinder2.0 [16] using inhibition readout, LL4 curve fitting, and ZIP model parameters.

## ACKNOWLEDGEMENTS

We thank Cindy Motaka for assistance in editing, Dr Olga Kharchenko for assistance with Figure 1, Dr Justin Patten (Davey Lab) for help with antiviral assay development and Dr. David Hesson for assistance in experimental design. J.F.D, M.K, J.C.P. Jr, N.S., D.B.S., and V.S.K. hold equity in Clear Creek Bio. M.K. and V.S.K are employees of Clear Creek Bio. V.G., D.C., J.F.D, N.S, J.C.P. Jr, and D.B.S. are paid consultants of Clear Creek Bio. D.B.S. holds equity in SAFI Biosolutions. Clear Creek Bio funded all work described in this article, including Sponsored Research Agreements with Boston University, MD Anderson Cancer Center and the University of Louisville.

